# Maloja: simple and scalable Snakemake workflow orchestration in the cloud

**DOI:** 10.1101/2024.06.28.601236

**Authors:** Joseph Giustizia, Wilson Hodgson, Cameron Andress, Satpal Bilkhu, James A Macklin, Tony Kess

**Affiliations:** Agriculture and Agri-food Canada, Ottawa, Canada; Northwest Atlantic Fisheries Centre, Fisheries and Oceans Canada, St. John’s, Newfoundland and Labrador, Canada

## Abstract

As sequencing technologies have matured, bioinformatics tasks have become more complex, computationally demanding, and data intensive. Workflow management software has been developed to aid in simplifying the replicable chaining of complex bioinformatics jobs, and cloud computing has emerged as a potential solution to the computational demands of this work. However, the capacity to effectively deploy these resources is limited by the expertise required to implement these solutions. Here, we develop Maloja, an easily deployed cloud workflow orchestrator. This tool interprets existing scientific workflows written in Snakemake and deploys them in appropriately scaled AWS cloud resources. We test the utility of this new toolset using previously published and custom built Snakemake workflows for ecological genomics tasks, revealing how this tool can facilitate the use of cloud resources without prior cloud architecture expertise.

## 1. Introduction

Over the past two decades, significant advances have been made in the availability and quality of molecular data. Initially, these advances have been due to rapid capacity increases in DNA sequencing methods, but other high-resolution approaches to measuring biological variation have also become feasible, leading to a new era of data-driven biology (Joyce and Pallson 2006; Ebrahim et al. 2016). These changes have been reflected in massive expansion of the volume of biological information retained in large digital archives. Scientific consortia have also developed around the utilization of these technologies to solve fundamental problems in biology and medicine (Zhou et al. 2022; Stobbe et al. 2022; *Anopheles gambiae* Genome Consortium 2020).

Research in these fields now routinely generates large datasets, often across multiple research teams, requiring enhanced infrastructure for data management and analysis among project participants (Schatz et al. 2023). As data volumes have increased, computational requirements have also expanded for research in omics fields (Stephens et al. 2015). These needs have been based on additional storage space for greater data volumes, increased compute capacity to analyze larger datasets, more complex algorithms and software pipelines and in-demand chip architectures such as graphics processing units (Nobile et al. 2017).

Cloud computing, the task-specific deployment of computing resources from a third party, has been utilized to solve some of these challenges, through the capacity to scale resources to large datasets and facilitate collaboration across different research teams. These services provide researchers with access to a larger pool of compute resources that are available within many institution-based clusters, as well as storage and sharing capacity for large genomic datasets (Langmead & Nallore 2018). Cloud-based toolsets have now been developed for automating components of genomics data analysis and processing (Nagasaki et al. 2023), single cell sequencing (Li et al. 2020) and proteomics (Muth et al. 2023), among many other bioinformatics tasks.

These changes in the hardware and environment for analysis of large omic datasets have been accompanied by the growing complexity of analysis toolsets. Researchers currently face challenges in managing complex software environments and dependencies, chaining together these tools, and sharing them reproducibly (Shade & Teal 2015). To meet this demand, scientific workflow management software such as NextFlow, SnakeMake, Common WorkFlow Language (CWL), and Workflow Description Language (WDL) have emerged as a high-level solution to these problems, and several workflow management frameworks have been developed to handle these challenges in scientific computing (Reiter et al. 2021). These tools allow the automated deployment of complex pipelines involving multiple analysis steps. Software components and versions are often abstracted in these workflow managers using package management or containerization tools, enhancing both the portability and reproducibility of these methods (Wratten et al. 2021).

Although cloud computing and workflow management tools provide advances in compute capacity and complexity to enable scientific computing on large omic datasets, there remain difficulties to their implementation across projects. Both of these resources require time investments to build expertise and familiarity with the language, logic, and performance of selected cloud environments and workflow managers. Additionally, managing a cloud environment requires dedicated management of security, software migration, and budgeting. Cloud environments also differ substantially from traditional scientific computing environments, and vary significantly depending on the cloud service provider, imposing a barrier to use. Integration of workflow management tools with cloud environments often requires alterations to the workflow description itself, and the inclusion of cloud-specific details to orchestrate and deploy cloud resources (Mölder et al. 2021). These constraints can add additional barriers for researchers primarily focused on shifting existing pipelines or standalone tools to cloud environments. As such, our objective is to explore the feasibility of developing a scalable and user-friendly tool for bioinformatics and cloud computing, with the aim of streamlining workflows and improving accessibility for researchers in the field.

Here, we describe the successful proof of concept development and testing of Maloja (pronounced “Ma-lo-ya”, named for a snake-shaped weather phenomenon: https://en.wikipedia.org/wiki/Maloja_Wind), a tool to facilitate easy deployment and management of scientific workflows in the cloud. Maloja is designed to be similar to the experience of working in a standard Unix-based high performance computing environment, allowing adoption of cloud tools without extensive cloud experience. Maloja utilizes python-based Snakemake workflow management system (Köster et al. 2012) to allow direct portability of workflows into an automated cloud deployment environment. This user design choice allows managed workflows across different data types to be easily shifted to cloud resources without re-engineering for cloud environments. We test this toolset in the AWS cloud environment using environmental DNA and bacterial genome assembly Snakemake workflows, highlighting the capacity of Maloja to enable cloud computing in ecological genomic research.

## 2. Overview and Architecture

Maloja is a Python-based command line utility that takes Snakemake workflows as input to create the appropriate cloud computing infrastructure (virtual machines, storage, networking, etc.) for running those workflows in an AWS environment (Figure 1). Cloud deployment of Snakemake workflows is performed with a simple python command line interface, obviating the need for tailoring code around individual workflows. To remove the barrier of cloud architecture design, Maloja instead auto-scales the required compute resources to user specified memory requirements, rather than requiring explicit virtual machine (VM) specifications (Figure 1).

**Figure 1.**
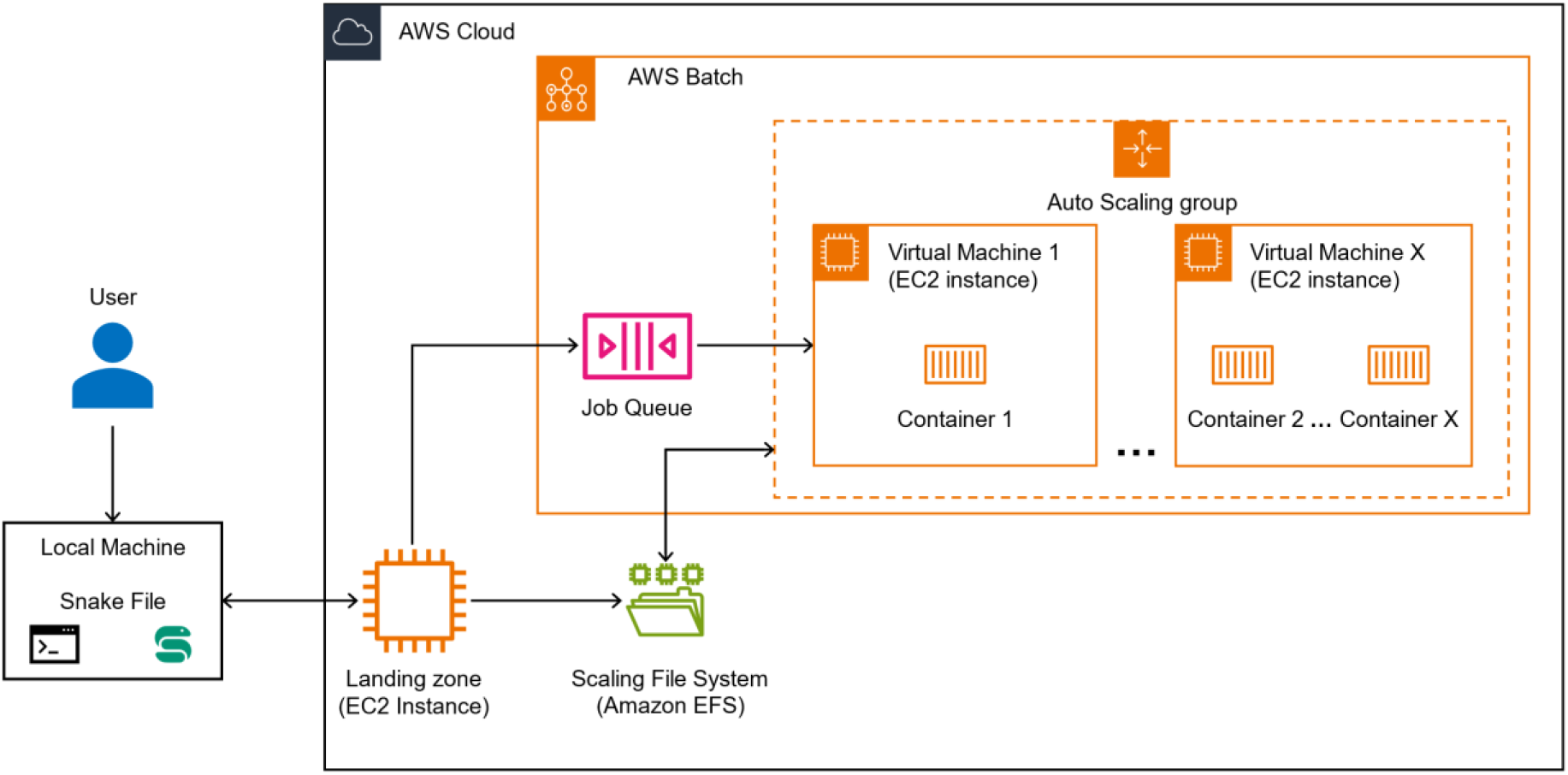
Maloja architecture overview. The user starts by engaging with their local computing environment to set up their pipeline definition. Then the pipeline is sent to the AWS Cloud landing zone to begin computation

These abstractions away from cloud engineering and architecture are achieved through the use of templating languages, which provide a structure in which variables are input to specify a desired task. This definition is specified only once, increasing standardization and reducing duplication of effort. Templating languages are the foundation of Maloja’s operation and are used to define both the analysis workflow, via SnakeMake, and to define the cloud infrastructure via AWS CloudFormation.

For users, Snakemake serves as the primary input mechanism, allowing researchers to describe computational tasks, dependencies, and resource requirements in a structured manner. For deployment of infrastructure, AWS CloudFormation templates derived from the resource specifications in Snakemake workflows are used to simplify the management of AWS resources by enabling the declaration of desired cloud infrastructure for pipeline execution within AWS environments. During pipeline initiation, Maloja also automatically creates a landing zone, from which both compute and storage instances are deployed, similar to the head node used for launching compute jobs in scientific computing clusters. Data storage during computation is managed in a scalable file system. Once analysis is triggered from the landing zone, an AWS batch instance is created which includes a job queue and an automatically-scaling set of virtual machines on which the analysis is run from within containers. AWS Batch efficiently streamlines the scheduling and launching of batch computing tasks. Upon completion of the pipeline, the results will be stored, and available for retrieval, in the scaling file system.

## 3. Testing

To test the utility of the Maloja cloud workflow orchestration tool, we tested two Snakemake pipelines: a recently published environmental DNA sequencing pipeline Metaworks 1.12.0 (Porter & Hajibabaei 2022), and a currently unpublished custom Snakemake pipeline developed for bacterial genome assembly. MetaWorks is a pipeline used in the automated analysis of environmental DNA datasets for species identification from environmental samples. For this pipeline, we used the test dataset provided in Metaworks, reflecting a COI MiSeq sequence dataset of 200MB size, with default parameters. The custom bacterial genome assembly pipeline was comprised of fastqc (Andrews 2010), multiqc (Ewels et al. 2016), bbduk (sourceforge.net/projects/bbmap/), shovill (https://github.com/tseemann/shovill), and prokka (Seeman 2014). We tested this assembly pipeline on four readsets of *Eschericihia coli* sequence data obtained from the Sequence Read Archive (SRR24437709, SRR24437710, SRR24437713, SRR24437715).

The total time required to deploy the infrastructure and run Metaworks on the test data was 1 hour 55 minutes. The run time for the Metaworks analysis with the provisioned architecture completed in 29 minutes and the cost was $2.20 USD. For the bacterial genome assembly pipeline, total run and infrastructure deployment time was: 2 hours 27 minutes, run time of the pipeline itself was 27 minutes, and the cost was $4.73 USD. See Table 1 for the cost and resource usage breakdown per pipeline.

**Table 1.**
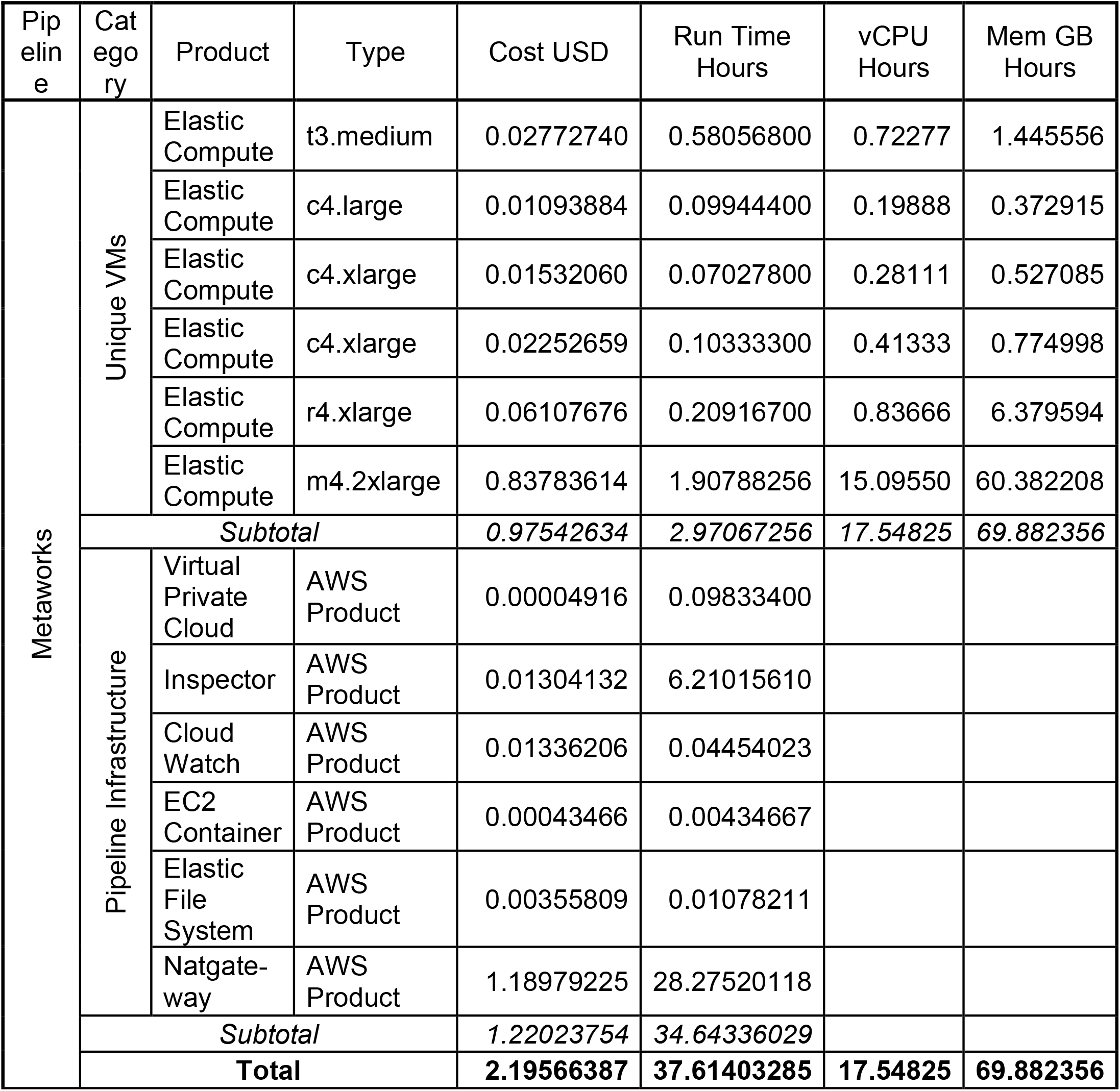
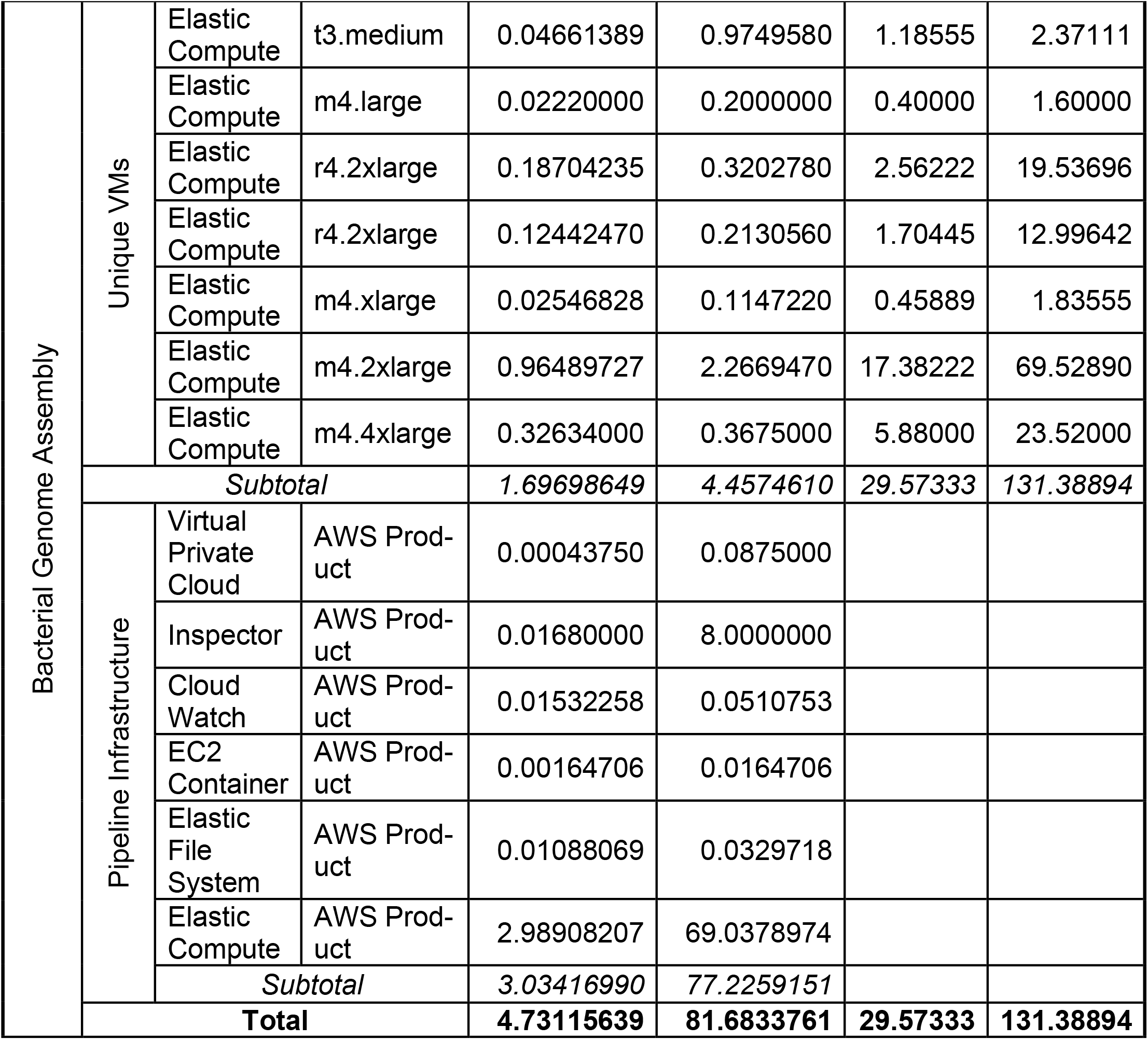
Amazon resources and associated summed costs per tested pipeline. All numerical values in this table are a summation of their categories. Note: these costs account only for AWS resource creation and usage. Environment idle time and any other potential costs are not reflected in this table

## 4. Discussion

Bioinformatic data analysis in the cloud is rapidly evolving, with a growing diversity of tools and over 150 workflow managers currently in use and under development (Wratten et al. 2021). [Across this diversity of methods, approaches differ in implementation details which prioritize different elements of user experience, automation, and interface choices](https://www.nature.com/articles/s41592-021-01254-9).

Here we have designed a cloud workflow tool that prioritizes familiarity by modelling the user experience on job scheduling tools that are commonly found in traditional scientific computing environments with a job scheduling system, such as SLURM (https://slurm.schedmd.com/documentation.html). We reduce the time investment associated with translating pipelines to cloud workflows by allowing provisioning of cloud resources to be handled without knowledge of specific cloud provider products offerings, instead simply specifying software and memory requirements for workflows.

We believe that this approach will aid in workflow management and cloud adoption by researchers not extensively familiar with both of these toolsets, and will reduce both time and difficulty associated with moving scientific computing into cloud environments. We have also designed this system with flexibility of input pipelines in mind, demonstrated this capacity by using both published and custom-built pipelines, and anticipated that pipelines respect best practices (e.g. Roach et al. 2022).

The recommended deployment approach by Snakemake for AWS is the use of Tibanna (Lee et al. 2019), an additional tool that can read pipelines in CWL, WDL, and Snakemake and run them on AWS. Maloja and Tibanna serve the same purpose overall of running snakemake workflows on AWS cloud environments, but differ in details of the design and implementation. One of the key differences between these tools is that Maloja uses Amazon’s Batch service, AWS Batch, which offers additional methods to manage autoscaling compute resources. Batch manages the compute through the definition of resources that are similar in function to the components of a cluster. These resources include Job definitions, queues for requesting jobs, and compute environments for executing these jobs. The compute environments in particular give an administrator of a Maloja deployment the ability to enforce limitations to the available compute resources for users. With respect to cost management, both Tibanna and Maloja can give overviews of the costs of running individual pipelines, but Maloja uniquely allows the constraining of allocated resources. Additionally, Maloja uses AWS Batch autoscaling to automatically increase and decrease the size and number of EC2 instances within a compute environment. Tibanna can use AWS to scale up the number of instances as well, but to the best of our knowledge this feature is only available to workflows defined in CWL and WDL, not to Snakemake native workflows. This means that Maloja is currently capable of saving costs on pipelines without requiring a conversion to WDL or CWL. Using AWS Batch has been an advantage to Maloja in improving both usability, and cost efficiency when running snakemake pipelines.

We demonstrate the deployment of both previously published (Metaworks) and custom built (bacterial genome assembly) pipelines in an AWS cloud environment facilitated by Maloja. Utilization of Maloja may facilitate user-friendly cloud computing for the analysis of bacterial genomes and environmental DNA, key use cases in ecological genomics that have been used for pathogen monitoring (Bass et al. 2022), ecosystem characterization (Crowley et al. 2024), and assessment of ecological restoration measures (Yan et al. 2018). The results from this proof of concept could be useful in enabling compute for ecological genomics applications requiring large compute and rapid turnaround time for characterization of microbial or ecological information, for example novel disease outbreaks or large scale environmental disasters such as oil spills. This approach can also be extended to other Snakemake workflows beyond those tested here, allowing rapid access to cloud-scale reproducible workflows across omic methods.

## Code availability

Source code and documentation for Maloja is available at https://github.com/GRDI-GenARCC/Maloja

## Acknowledgements

We thank the Genomics Research and Development Initiative for financial support through the Genomic Adaptation and Resilience to Climate Change (GenARCC) project and Shared Services Canada for access to compute resources and logistic support. Helpful comments on an earlier version of the manuscript were provided by Jackson Eyres and Etienne Low-Décarie.

## References

Andrews, S. (2010). FastQC: A quality control tool for high throughput sequence data. http://www.bioinformatics.babraham.ac.uk/projects/fastqc.

The Anopheles gambiae 1000 Genomes Consortium. (2020). Genome Variation and Population Structure among 1142 Mosquitoes of the African Malaria Vector Species Anopheles Gambiae and Anopheles Coluzzii. Genome Res, 30(10):1533–1546. 10.1101/gr.262790.120.

Bass D, Christison KW, Stentiford GD, Cook LSJ, Hartikainen H. (2023). Environmental DNA/RNA for pathogen and parasite detection, surveillance, and ecology. Trends Parasitol, 39(4):285–304. 10.1016/j.pt.2022.12.010.

Bushnell, B. (n.d.). BBMap. Retrieved from https://sourceforge.net/projects/bbmap/

Ebrahim, A., Brunk, E., Tan, J., et al. (2016). Multi-omic data integration enables discovery of hidden biological regularities. Nat Commun, 7:13091. 10.1038/ncomms13091.

Ewels, P., Magnusson, M., Lundin, S., & Käller, M. (2016). MultiQC: summarize analysis results for multiple tools and samples in a single report. Bioinformatics, 32(19), 3047–3048. 10.1093/bioinformatics/btw354

Joyce, A., Palsson, B. (2006). The model organism as a system: integrating ‘omics’ data sets. Nat Rev Mol Cell Biol, 7:198–210. 10.1038/nrm1857.

Köster, J., Rahmann, S. (2012). Snakemake—a Scalable Bioinformatics Workflow Engine. Bioinformatics, 28(19):2520–2522. 10.1093/bioinformatics/bts480.

Langmead, B., Nellore, A. (2018). Cloud computing for genomic data analysis and collaboration. Nature Reviews Genetetics, 19:208–219. 10.1038/nrg.2017.113.

Lee, S., Johnson, J., Vitzthum, C., Kirli, K., Alver, B. H., & Park, P. J. (2019). Tibanna: software for scalable execution of portable pipelines on the cloud. Bioinformatics, 35(21), 4424–4426. 10.1093/bioinformatics/btz379.

Li, B., Gould, J., Yang, Y., et al. (2020). Cumulus provides cloud-based data analysis for large-scale single-cell and single-nucleus RNA-seq. Nature Methods, 17:793–798. 10.1038/s41592-020-0905-x.

Mölder F, Jablonski KP, Letcher B, Hall MB, Tomkins-Tinch CH, Sochat V, Forster J, Lee S, Twardziok SO, Kanitz A, Wilm A, Holtgrewe M, Rahmann S, Nahnsen S, Köster J. (2021). Sustainable data analysis with Snakemake. F1000Res, 10:33. 10.12688/f1000research.29032.2.

Muth, T., Peters, J., Blackburn, J., Rapp, E., & Martens, L. (2013). ProteoCloud: a full-featured open source proteomics cloud computing pipeline. Journal of Proteomics, 88, 104–108. 10.1016/j.jprot.2012.12.026.

Nagasaki, M., Sekiya, Y., Asakura, A., et al. (2023). Design and implementation of a hybrid cloud system for large-scale human genomic research. Human Genome Variation, 10:6. 10.1038/s41439-023-00231-2.

Nobile MS, Cazzaniga P, Tangherloni A, Besozzi D. (2017). Graphics processing units in bioinformatics, computational biology and systems biology. Brief Bioinform, 18(5):870–885. 10.1093/bib/bbw058.

Porter TM, Hajibabaei M. (2022). MetaWorks: A flexible, scalable bioinformatic pipeline for high-throughput multi-marker biodiversity assessments. PLoS ONE, 17(9):e0274260. 10.1371/journal.pone.0274260.

Reiter, T., Brooks, P. T., Irber, L., Joslin, S. E. K., Reid, C. M., Scott, C., Brown, C. T., Pierce-Ward, N. T. (2021). Streamlining data-intensive biology with workflow systems. GigaScience, 10(1):giaa140. 10.1093/gigascience/giaa140.

Roach MJ, Pierce-Ward NT, Suchecki R, Mallawaarachchi V, Papudeshi B, Handley SA, et al. (2022). Ten simple rules and a template for creating workflows-as-applications. PLoS Comput Biol, 18(12):e1010705. 10.1371/journal.pcbi.1010705.

Schatz, M. C., Philippakis, A. A., Afgan, E., Banks, E., Carey, V. J., Carroll, R. J., Culotti, A., … & Walker, J. (2022). Inverting the Model of Genomics Data Sharing with the NHGRI Genomic Data Science Analysis, Visualization, and Informatics Lab-Space. Cell Genomics, 2(1), 100085. 10.1016/j.xgen.2021.100085.

Seemann, T. (2014). Prokka: rapid prokaryotic genome annotation. Bioinformatics, 30(14), 2068–2069. 10.1093/bioinformatics/btu153

Shade, A., & Teal, T. K. (2015). Computing Workflows for Biologists: A Roadmap. PLOS Biology, 13(11), e1002303. 10.1371/journal.pbio.1002303.

Stephens, Z. D., Lee, S. Y., Faghri, F., Campbell, R. H., Zhai, C., Efron, M. J., Iyer, R., Schatz, M. C., Sinha, S., & Robinson, G. E. (2015). Big data: astronomical or genomical?. PLoS Biology, 13(7), e1002195.

Wratten, L., Wilm, A., & Göke, J. (2021). Reproducible, Scalable, and Shareable Analysis Pipelines with Bioinformatics Workflow Managers. Nature Methods, 18(10), 1161–1168. 10.1038/s41592-021-01254-9.

Yan, D., Mills, J. G., Gellie, N. J., Bissett, A., Lowe, A. J., & Breed, M. F. (2018). High-throughput eDNA monitoring of fungi to track functional recovery in ecological restoration. Biological Conservation, 217, 113–120. 10.1016/j.biocon.2017.10.035.

